# Chronic Alcohol Consumption Alters Home-Cage Behaviors and Responses to Ethologically Relevant Predator Tasks in Mice

**DOI:** 10.1101/2022.02.04.479122

**Authors:** Sofia Neira, Leslie A. Hassanein, Christina M. Stanhope, Michelle C. Buccini, Shannon L. D’Ambrosio, Meghan E. Flanigan, Harold L. Haun, Kristen M. Boyt, Jaideep S. Bains, Thomas L. Kash

## Abstract

Alcohol use disorders (AUD) are the most prevalent substance use disorders worldwide. Considering recent reports indicating an increase in alcohol use particularly in females, it is vital to understand how alcohol history impacts behavior. Animal model research on withdrawal-associated affective states tends to focus on males, forced alcohol paradigms, and a few traditional anxiety/stress tests. While this has been essential, heavy alcohol use triggers adverse withdrawal-related affective states that can influence how people respond to a large variety of life events and stressors. To this end, we show that behaviors in the home-cage, open field, looming disc, and robogator predator threat task, which vary in task demand and intensity, are altered in mice with a history of voluntary alcohol consumption. In alcohol-exposed males, behaviors in the home cage, a low anxiety baseline environment, suggest increased vigilance/exploration. However, in the open field and robogator task, which induce heightened arousal and task demands, a more hesitant/avoidant phenotype is seen. Female alcohol mice show no behavioral alterations in the home cage and open field test, however, in the looming disc task, which mimics an overhead advancing predator and forces a behavioral choice, we see greater escape responses compared to water controls, indicative of active stress coping behaviors. This suggests females may begin to show alcohol-induced alterations as task demands increase. To date, few drugs have advanced past clinical trials for the treatment of AUD, and those that have are predominately used in life-threatening situations only. No treatments exist for ameliorating negative withdrawal related states, which could aid in harm reduction related to heavy alcohol use. Understanding how withdrawal alters a variety of behavioral responses that are linked to stress coping can widen our understanding of alcohol abuse and lead us closer to better therapeutics to help individuals with AUD.

## Introduction

Alcohol Use Disorders (AUD) affect approximately 14.5 million people in the United States (SAMHSA, 2019). The stress associated with negative affective states during withdrawal is thought to play a role in the resumption of alcohol use and thus in the perpetuance of AUDs (Becker, 2012; Blaine et al., 2019; Blaine and Sinha, 2017; Heilig et al., 2010; Magrys et al., 2013). This ‘dark side of addiction’ (Koob and Le Moal, 2005) has led to extensive work using animal models on how stress drives drinking patterns with a goal of finding targets for treatments that reduce alcohol consumption (Deal et al., 2018; Haun et al., 2022; Koob and Mason, 2016; Lê et al., 2011; Newman et al., 2018). In comparison, relatively little is known about how a history of alcohol use affects stress responses despite the growing evidence in human studies on alcohol-associated alterations to Hypothalamic Pituitary Adrenal (HPA) axis physiological and behavioral responses to stress (Blaine et al., 2019; Blaine and Sinha, 2017; Metzger et al., 2018; Sinha et al., 2009). All organisms respond to stress by marshaling physiological responses (i.e., changes in hormones, heart rate, blood pressure, respiration) and context-appropriate behaviors (i.e flight/freeze response to acute stress, exploration in food-seeking, self-grooming in rodents for de-arousal after a stressor) that depend on the stressor, arousal level, and the species (Andrew, 1974; Bracha, 2004; Pankevich and Bale, 2008; Veloso et al., 2016). The behavioral impact of alcohol withdrawal on stress, both generalized and stressor specific, has not been heavily explored.

Recent preclinical studies investigating negative affective states during withdrawal from voluntary, recurring alcohol consumption show behavioral effects with considerable variability in findings using traditional stress/anxiety tests (Bloch et al., 2020; Kliethermes, 2005). This could be due to differences in drinking paradigms used across experiments that result in distinct intake patterns and overall consumption. Recent analyses, however, have questioned the validity of commonly used behavioral approaches for assessing negative emotional behaviors in rodents, arguing that their reliance on anthropomorphism undermines their relevance and consistency (Kliethermes, 2005; Lezak et al., 2017). In step with this criticism, elements of behavioral research have moved towards using etho-experimental approaches to behavioral assessment which might prove useful in investigating stress and anxiety-like behaviors in alcohol withdrawal. This includes detailed analysis of spontaneous, natural behavior patterns (i.e. grooming, rearing, digging, locomotion) and ethologically and species relevant behavioral tasks, such as those that drive behavioral choices in the presence of threats or hunger (Blanchard et al., 1993; Blanchard and Blanchard, 1990; Lezak et al., 2017; Richter, 2020). Here we focus on spontaneous home-cage behavior in mice (Füzesi et al., 2016), as well as two main ethologically and species-relevant tests for examining innate defensive behavior:the looming disc (Daviu et al., 2020), and the robogator-simulated predator task (Choi and Kim, 2010), to determine how stress coping and stress response behaviors are altered in mice during forced abstinence from six weeks of two bottle choice voluntary alcohol consumption.

## Materials and Methods

### Animals

Group housed male and female C57BL/6J mice (Jackson Laboratories, Bar Harbor, ME) were delivered at 6-8 weeks of age and maintained on a 12-h reverse dark/light cycle (light off at 7:00am). After at least one week of habituation to the animal facility, mice were single housed in polycarbonate (GM500, Tecniplast, Italy) Plexiglas cages, given an additional 5-7 days to adjust to the single housing conditions, and assigned to the water control group or two bottle choice alcohol access group. Subjects were matched for age and weight (N = 98 males:N = 73 females). Unless otherwise specified, mice were given *ad libitum* access to water and Isopro RMH 3000 chow (LabDiet, St. Louis, MO). Behavioral experiments were conducted during the dark cycle and with the room lights off, and males and females were run separately. The Institutional Animal Care and Use Committee at UNC Chapel Hill approved all experimental procedures, which were performed in accordance with the NIH Guide for the Care and Use of Laboratory Animals.

### Two Bottle Choice Alcohol Access

For six weeks, alcohol mice were given 24-hour access to water and a 20% (w/v) alcohol solution (water and 95% ethanol:Pharmaco-AAPER, Brookfield, Connecticut) on Mondays, Wednesdays, and Fridays for a total of 18 alcohol drinking sessions. Both water and alcohol bottles were weighed before and after each 24-hour drinking session, and the averaged value from an empty ‘drip’ cage was subtracted from the overall drinking value. The final g/kg consumption was determined with this formula:((alcohol consumed in ml) * 0.206) / (weight of mouse in g /1000). All mice, including the water access control groups, had unrestricted access to water throughout the six-week access paradigm. Ten mice used in Experiment 2 only received 16-17 drinking sessions due to COVID-19 related experimental issues.

### Experiment 1:Validation of alcohol drinking paradigm

Four hours into the final, 18th drinking session, mice were anesthetized in a small isoflurane chamber, removed from the animal facility, and sacrificed. Trunk blood was immediately collected in a Sarstedt Inc heparinized microvette CB300 tube (Fisher Scientific, Waltham, MA). Blood samples were centrifuged at 4 °C for 10 min at 3000 RPM, and the plasma was separated. Blood ethanol content (B.E.C.; mg/dL) was analyzed immediately with a 5-µL plasma sample using a Model AM1 Alcohol Analyser (Analox Instruments Ltd. Lunenburg, Massachusetts).

### Experiment 2:Effects of alcohol on home-cage and open field behaviors

24-36 hours after the final water or alcohol drinking session, mice were placed on a cart, covered, and transferred to the behavioral testing room. After a minimum of 40-mins acclimation to the room, mice were either left undisturbed in their home-cage (No-Stress) or restrained for 2-hours (Stress). Immediately following the No-Stress or Stress conditions, mice were subjected to Home Cage and Open Field tests. Using the free event logging software BORIS (Friard and Gamba, 2016), a blinded individual scored grooming, rearing, and digging behaviors. Tracking and locomotor activity were analyzed using EthoVision (Noldus Information Technologies, Wageningen, Netherlands) and DeepLabCut (DLC) and Simple Behavioral Analysis (SimBA) machine learning based models (DLC:https://github.com/DeepLabCut/DLCutils; SimBA:https://github.com/sgoldenlab/simba).

#### Restraint Stress

Mice were scruffed and placed in a well-ventilated 50mL conical centrifuge tube (Thermo Scientific, Fisherbrand, Waltham, Massachusetts) for two hours in a dark enclosed space. Crumpled paper towels were placed in between the mouse and the cap of the tube to further restrict movement.

#### Home-Cage

The home-cage was cleared of all items but bedding. No-Stress mice were picked up and placed back in the home-cage and Stress mice were removed from restraint tube and placed back in their home-cage. The cages were then moved to a low light (∼60 lux) enclosed space with a webcam placed above to film home-cage behaviors for 15 minutes.

#### Open Field

Immediately following the home-cage behavioral test, mice were placed in a white Plexiglas open field (50 × 50 × 25 cm) and allowed to explore the arena for 5 minutes. Light levels in the center of the open field were approximately 300 lux.

#### DeepLabCut and SimBA

This behavioral experiment was conducted by multiple researchers in separate cohorts. The final male cohort and the female cohorts were filmed using higher resolution cameras (C930e Webcam, Logitech, Lausanne, Switzerland) compatible with automatic behavioral analysis pipelines using machine learning. DLC (Mathis et al., 2018), a free open access machine learning program for behavioral pose estimation, was used to track the mouse across the home-cage. This tracking information was then used to create behavioral classifiers for the most displayed home-cage behaviors using SimBA (Nilsson et al., 2020), a separate, open source program for identifying specific behaviors. We followed the extensive existing GitHub documentation for setup and function of these programs and developed our own tracking DLC project and 3 models for rearing, digging, and grooming in SimBA using the DLC tracking information. The ethovision tracking information was compared to DLC/SimBA tracking, and the data derived from the SimBA models was compared to the blinded hand-scoring of the behaviors in a subset of videos from experiment two to confirm consistent results.

### Experiment 3: Effects of alcohol on the looming disc task

Twice during the final week of water or alcohol access, mice were placed on a covered cart and brought upstairs to the testing room on alcohol “off” days. After a minimum of 40-min acclimation to the testing room in the home-cage, mice were habituated to the looming disc environment (a 41 × 19 × 20.5 cm mirrored floor plastic arena with a hut (13 × 12 × 10cm) in one corner) for 20-minutes per day. A computer monitor (22er 21.5-inch LED Backlit, HP inc., Palo Alto, California) was placed above the arena with a light grey background that produced an ambient light of approximately 170 lux. On test day, 6-8 hours after the final water or alcohol drinking session, mice were once again brought to the testing room.

#### Looming Disc

After a minimum of 40-min acclimation to the room, mice were subjected to the looming disc task (Daviu et al., 2020). Mice were given 3 minutes in the looming disc arena for habituation. Following that period, a loom was triggered when the mouse was not in the hut or heading towards the hut. A loom was then triggered an additional 4 times, with at least thirty seconds elapsing between each loom. The loom consists of a 2 cm black disc on a light grey background. The disc is first presented for 3 seconds, and then expands to its full 20cm size over 2 seconds. The 20cm disc stays on the screen for an additional 3 seconds and disappears. A blinded individual analyzed mouse behavior, with the criteria of escape entailing reaching the hut within the 8 second loom presentation and freeze being the absence of movement other than breathing for at least 1 second total during loom presentation.

### Experiment 4:Effects of alcohol on the robogator predator task

Based on previous experiments using a moving predator-like Lego robot, or robogator, (Choi and Kim, 2010), starting 3 days after the final water or alcohol access session, male mice were given chocolate sucrose pellets (Dustless Precision Pellets, Bio-Serv, Flemington, NJ) daily for 4 days in their home-cage to reduce neophobia. We began this task seven days after the final drinking session. Mice were first food deprived for approximately 18 hours prior to habituation day and were kept at 85% or more of their original weight throughout the entire test period by food restriction. Food was given immediately after the mouse finished each task. Criteria for moving past day 4 of baseline training to test day was to retrieve the three pellets by 300 seconds each. One mouse from the alcohol access group was excluded due to health concerns and one water mouse was excluded for failure to reach the criteria for entering test day. This experiment was conducted in female mice using a different approach with higher stress conditions; thus, only male data is included in this manuscript.

#### Robogator Test

Habituation –To associate the arena with food foraging and to habituate and introduce the mice to the nest and field of the robogator arena (nest:8 × 19.5 × 24 inches, field:40 × 19.5 × 24 inches), mice were first allowed to explore the nest for 10 minutes and then a gate was raised to give access to the field. Mice were then given unrestricted access to the entire arena for an additional 20 minutes. Food pellets were dispersed randomly throughout the arena.

Baseline Training – For 4 days, mice were restricted in the nest for 1 minute and trained to retrieve a pellet from each of the 30cm, 60cm, and 90cm distances from the nest. Test Day – Mice were restricted to the nest for 1 minute and then given access to the arena for 3 minutes to retrieve a pellet in the presence of the robogator (LEGO MINDSTORMS EV3, The Lego Group, Billund, Denmark) robot, which moved forward, snapped its “jaws” 3 times, and moved back when the mouse was approximately 30cm or closer to the food pellet. All mice began with the first trial – where the pellet was placed 30cm from the nest. Subsequent trials depended on the success of the mouse in retrieving a pellet. If successful, the pellet was moved 15cm further away from the nest (45cm, 60cm, 75cm, 90cm). If a mouse failed, the trial began with the pellet 15cm closer to the nest (15cm, 0cm). Trials continued until the maximum foraging distance (MFD) was determined. The MFD was the final distance from the nest entrance (1cm, 15cm, 30cm, 60cm, 75cm, or 90cm) where each mouse was successful in obtaining a pellet in less than 3 minutes. The pellet was moved back to its original spot in the center of the field if it was accidentally moved by the mouse as it responded to the robogator.

#### Statistics

When appropriate, student’s two-tailed t-tests, mixed-effects model (in cases where data points are missing), one-way repeated measures (RM) ANOVAs, two-way RM ANOVAs, or two-way ANOVAs were conducted. Tukey’s corrected multiple comparison post-hoc tests were performed when applicable. To explore the relationship between two variables, simple linear regressions were performed. Non-parametric data was compared using a Fisher’s exact test. We also conducted a survival analysis, an epidemiological statistical method that can be used to understand animal behavior (Asher et al., 2017), on the robogator MFD. A MFD of 1cm -75cm was counted as a ‘death’, while mice foraging up to the final distance of 90cm were counted as ‘surviving’. A p-value of less than 0.05 was considered significant and data is shown as individual values with mean and standard error of the mean. When individual values are not shown, data is shown as mean and 95% confidence interval. Statistical analyses were conducted using Prism 9.2.0 (GraphPad Software Inc, San Diego, CA).

## Results

### Experiment 1

Our interest lies in understanding how voluntary alcohol intake affects subsequent behaviors during withdrawal. Much of the field focuses on forced alcohol models to induce high B.E.C.s, high withdrawal symptoms, and low variability. While this is critical to understanding aspects of severe AUD, many people with AUD have milder consumption patterns and symptomology, and forced alcohol paradigms may be overrepresenting those more extreme cases (Holleran and Winder, 2017). Therefore, we first explored intake and preference patterns in male and female mice in the 6-week two-bottle choice paradigm (Fig 1A) and tested whether this paradigm would induce pharmacologically relevant B.E.C.s at the standard 4hr timepoint used in drinking in the dark (DID) models. We first pooled together all of the daily intake and preference values for every mouse (excluding the 18^th^ session of experiment 1 mice that only drank for 4hrs; N= 52 males and 43 females) and found a significant difference in intake (Fig 1B:

**Figure 1:**
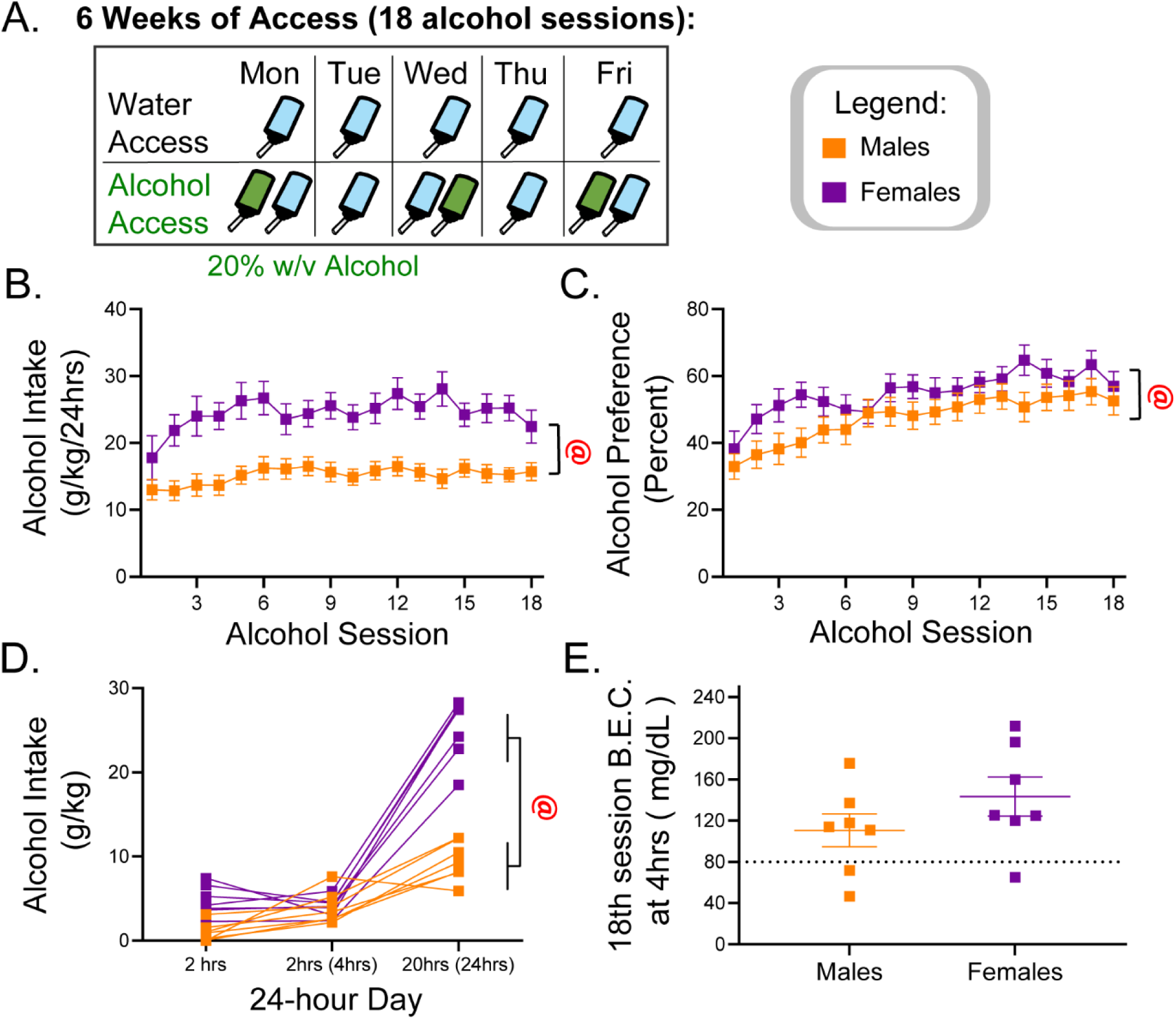
6-week two-bottle choice alcohol (20% w/v) access with forced 24-48hr withdrawals leads to distinct drinking patters in male and female mice. (A) Two-bottle choice schedule. (B) Daily intake (g/kg/24hrs; Mixed-effects model, Sex [F(1,95) = 125.5, p<.001]) and (C) preference (percent; Mixed-effects model, Sex [F(1,95) = 12.44, p<.001]) over the 18 alcohol sessions. Experiment 1 (D) individual mouse alcohol intake patterns over a 24hr session and (E) Blood alcohol content (mg/dL) four hours into the final, 18^th^ alcohol session.

Mixed-effects model, Sex [F(1,95) = 125.5, p<.001]) and preference (Fig 1C:Mixed-effects model, Sex [F(1,95) = 12.44, p<.001]) between male and female mice. Intake and preference also differed across the 6-weeks, with females showing greater escalation in alcohol intake (Fig 1B:Mixed-effects model, Time [F (9.094, 831.8) = 9.131, p<.001]; Time x Sex [F(17,1555) = 3.229.5, p<.001]), and both males and females showing a clear increase in preference for alcohol across time (Fig 1C:Time [F (11.95, 1127) = 34.70, p<.001]; Time x Sex [F(17,1602) = 3.150, p<.001]). In a subset of these mice that were used in experiment 1, we observed alcohol intake across a 24-hr session in the final week of alcohol exposure and found that males and females showed variability in drinking between subjects as well as distinct patterns of intake during the day (Fig 1D:Two-way RM ANOVA, Sex [F(1,12) = 60.84, p<.001]; Time [F(1.517,18.20) = 274.7, p<.001]; Time x Sex [F(2,24) = 70.90, p<.001]; Subject [F(12,24) = 2.232, p<.05]). Despite these differences in drinking patterns, most mice achieved binge drinking above 80mg/dl B.E.C. levels in a 4hr alcohol session on the final day of access (Fig. 1E). Ultimately, due to the highly distinct patterns of alcohol intake between sexes, males and females were analyzed separately in subsequent experiments.

### Experiment 2

We next sought to determine how behaviors in the home cage and open-field arena were impacted by withdrawal from alcohol in combination with acute stress exposure (Fig 2A). Spontaneous home cage behaviors, such as grooming, digging, and rearing are altered after a stress exposure (Füzesi et al., 2016) and may reflect heightened anxiety-like behavior. Thus, we evaluated these behaviors at baseline or after a single 2hr restraint stress in water and alcohol access mice. Restraint stress is known to potentiate anxiety-like behavior observed during withdrawal from alcohol (Breese et al., 2005, 2004) and thus was selected for these experiments.

**Figure 2:**
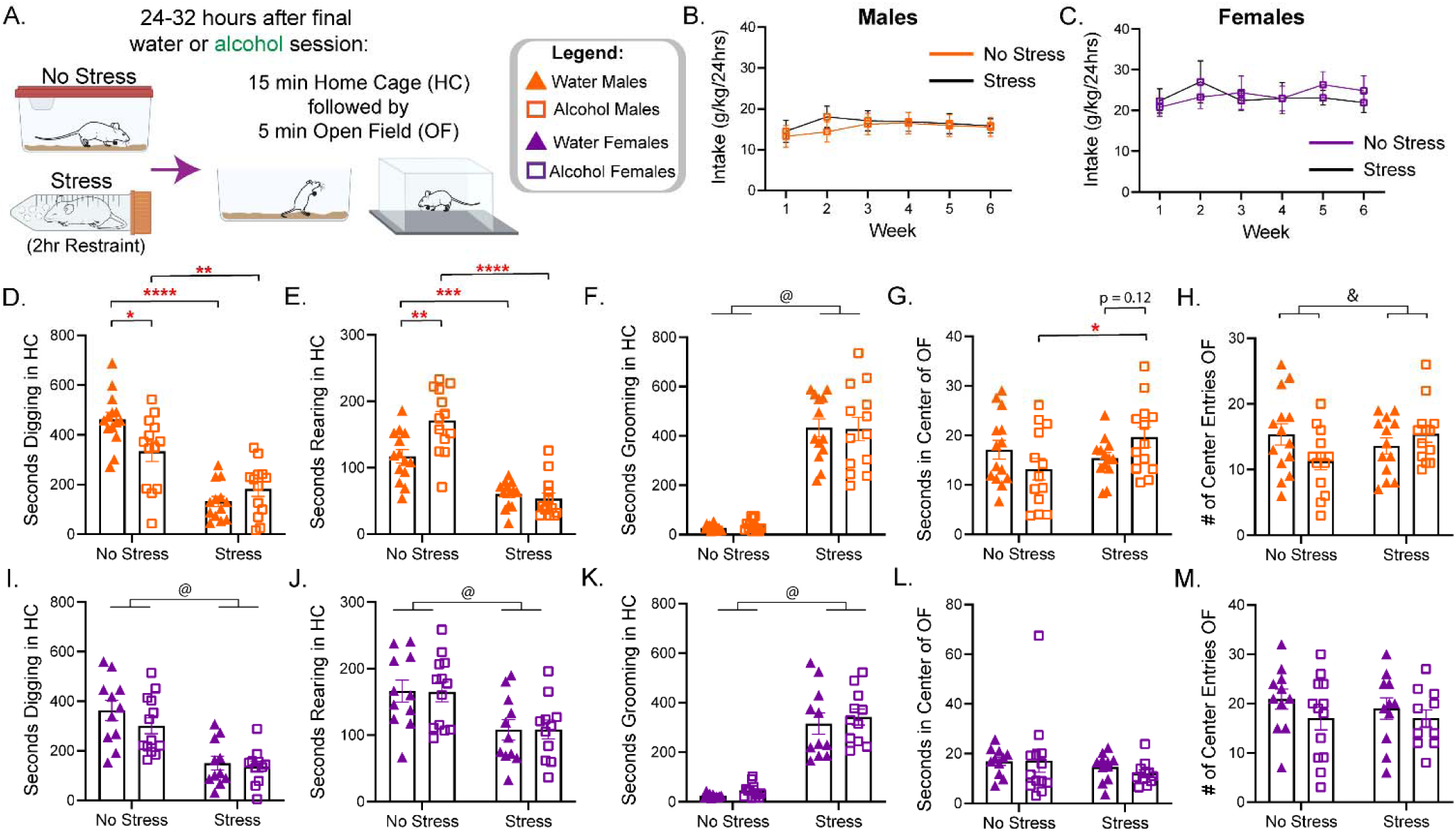
Acute alcohol withdrawal from long term access drives alterations in Home-Cage and Open Field behaviors in male mice only while acute, 2hr restraint stress drives similar alterations in male and female mice regardless of alcohol history. (A) Experimental design. (B) Alcohol intake in male mice (Weekly averaged g/kg/24hrs; 2way RM ANOVA, Condition [F(1, 24) = 0.6818, n.s.]). (C) Alcohol intake in female mice (Weekly averaged g/kg/24hrs; 2way RM ANOVA, Condition [F(1, 22) = 0.1089, n.s.]). In Males, Home Cage test:(D) Seconds spent digging (2way ANOVA, No-Stress water vs alcohol p = 0.0211, No-Stress water vs Stress water p < 0.0001, No-Stress alcohol vs Stress alcohol p < 0.0001). (E) Seconds spent rearing (2way ANOVA, No-Stress water vs alcohol p = 0.0015), No-Stress water vs Stress water p = 0.0008, No-Stress alcohol vs Stress alcohol p < 0.0001). (F) Seconds spent grooming (two-way ANOVA, main effect of Stress [F(1, 49) = 182.1, p < 0.0001]). In males, Open Field test:(G) Seconds spent in the center of the open field (2way ANOVA, No-Stress alcohol vs Stress alcohol p = 0.018) and (H) number of center entries (2way ANOVA, drinking x stress interaction:p = 0.039). In females, Home Cage test:(I) Seconds spent digging (2way ANOVA, Main effect of Stress [F(1, 42) = 35.71, p<.001]) (J) Second spent rearing (2way ANOVA, Main effect of Stress [F(1, 42) = 14.01, p<.001]). (K) Second spent grooming (two-way ANOVA, main effect of Stress (Fig 2K [F(1, 42) = 127.3, p<.001]). In females, Open Field test:(L) Seconds in the center of the open field and (M) number of center entries. @ = Main effect of stress. & = Alcohol x Stress Interaction effect. Tukey’s Post hoc tests:* = p<.05, ** = p<.01, *** = p<0.001, **** = p<.0001.

Mice were separated into treatment conditions prior to testing and No-Stress and Stress alcohol male (Fig 2B:2way RM ANOVA, Condition [F(1, 24) = 0.6818, n.s.]) or female (Fig 2C:2way RM ANOVA, Condition [F(1, 22) = 0.1089, n.s.]) mice did not differ in intake over the six weeks of consumption. In males, for time spent digging (Fig 2D), a two-way ANOVA indicated a significant interaction between alcohol drinking and Stress [F(1, 49) = 8.542, p = 0.0052], as well as a main effect of Stress [F(1, 49) = 61.86, p < 0.0001], but not alcohol [F(1, 49) = 1.655, p = 0.2043]. Post hoc tests indicated that alcohol No-Stress mice spent less time digging than water No-Stress controls (p = 0.0211). Stress led to a large reduction in digging in both water and alcohol access mice (water No-Stress vs water Stress p < 0.0001; alcohol No-Stress vs alcohol Stress p = 0.0060). Rearing time (Fig 2E) was impacted by both alcohol consumption and stress (two-way ANOVA; main effect of alcohol [F(1, 49) = 5.708, p = 0.0208]; Stress [F(1, 49) = 78.49, p < 0.0001]; and an interaction [F(1, 49) = 9.617, p = 0.0032]). Post hoc tests indicated no differences between the stress conditions (p = 0.9589), while there was a significant increase in time spent rearing in No-Stress alcohol mice compared to No-Stress water mice (p = 0.0015). Stress caused a similar reduction in rearing time (water No-Stress vs water Stress p = 0.0008; alcohol No-Stress vs alcohol Stress p < 0.0001). As expected, grooming vastly increased following stress (two-way ANOVA, main effect of Stress F(1, 49) = 182.1, p < 0.0001], but there was no main effect of drinking condition or an interaction effect.

We next wanted to explore how alcohol and stress affect behaviors in a more widely used test to assess aspects of anxiety-like behavior, the open field. Two-way ANOVA analyses indicated no main effects and an interaction effect between alcohol and stress in two factors closely tied to anxiety, the number of center entries (Fig 2G, two-way ANOVA; Alcohol x Stress interaction [F(1, 49) = 4.5, p = 0.039]), and the amount of time spent in the center of the open field (Fig 2H, two-way ANOVA; Alcohol x Stress interaction [F(1, 49) = 4.846, p = 0.032]). Post-hoc tests indicated no differences in the center entries, while the time spent in the center was trending to significance between stress mice (alcohol Stress vs water Stress p = 0.12) and significantly different between No-Stress alcohol and Stress alcohol mice (p = 0.018). These effects were not likely due to differences in locomotion, as there were no differences in distance moved between the naïve water and alcohol groups in the home cage or open field tests. There was a reduction in distance moved in Stress mice in the home cage, however, this was likely due to the significant amount of time spent grooming (Supplementary Fig 1A-B).

Interestingly, in females, alcohol history alone did not drive any differences in home cage behaviors or the open field test, while restraint stress drove similar changes in both water and alcohol exposed mice that mirror those seen in stressed males (Fig 2D-M). Restraint stress drove a reduction in digging (Fig 2; Main effect of Stress [F(1, 42) = 35.71, p<.001]) and rearing (Fig. 2J; Main effect of Stress [F(1, 42) = 14.01, p<.001]) behaviors, while grooming drastically increased (Fig 2K; Main effect of Stress [F(1, 42) = 127.3, p<.001]). In the open field test, no differences were seen between the groups or the Stress condition in the time spent in the center (Fig 2L) or the center entries (Fig 2M). Female No-Stress mice did not differ in distance moved in the home cage or open field, but there was a reduction in the Stress condition mice in the home cage, likely due to the high amount of time spent grooming (Supplementary Fig 1C-D).

The home cage test indicated that in males, withdrawal from alcohol consumption alone led to alterations to spontaneous baseline behaviors in a familiar environment (Fig 2D-F). Due to the time-consuming aspect of hand scoring spontaneous behaviors, we explored the value of using DLC/SimBA machine learning for analyzing this data (Fig 3). We explored the relationship between Ethovision-sourced locomotion data and DLC/SimBA data, as well as the blinded hand-scoring of the three behaviors with the SimBA created models. Locomotion data was highly correlated, however, Ethovision had locomotion data approximately 3X higher than DLC/SimBA (Supplementary fig. 1E, simple linear regression [F(1, 13) = 699.3, p <0.0001]). The grooming (Fig 3A, simple linear regression R^2^ = 0.98, [F(1, 13) = 514.1, p <0.0001]) and rearing (Fig 3B, simple linear regression R^2^ = 0.93, [F(1, 13) = 182.7, p <0.0001]) SimBA models were highly correlated to hand scored data. Interestingly, the digging model did not match the hand scored data as closely as the other two (Fig 3C, simple linear regression R^2^ = 0.77 [F(1, 13) = 42.72, p <0.0001]), which might reflect the harder start/stop distinction of this behavior in hand-scoring. Overall, machine learning results are well matched to blind behavior scoring and can be used for a rapid, unbiased, and robust analysis of these types of behaviors.

**Figure 3:**
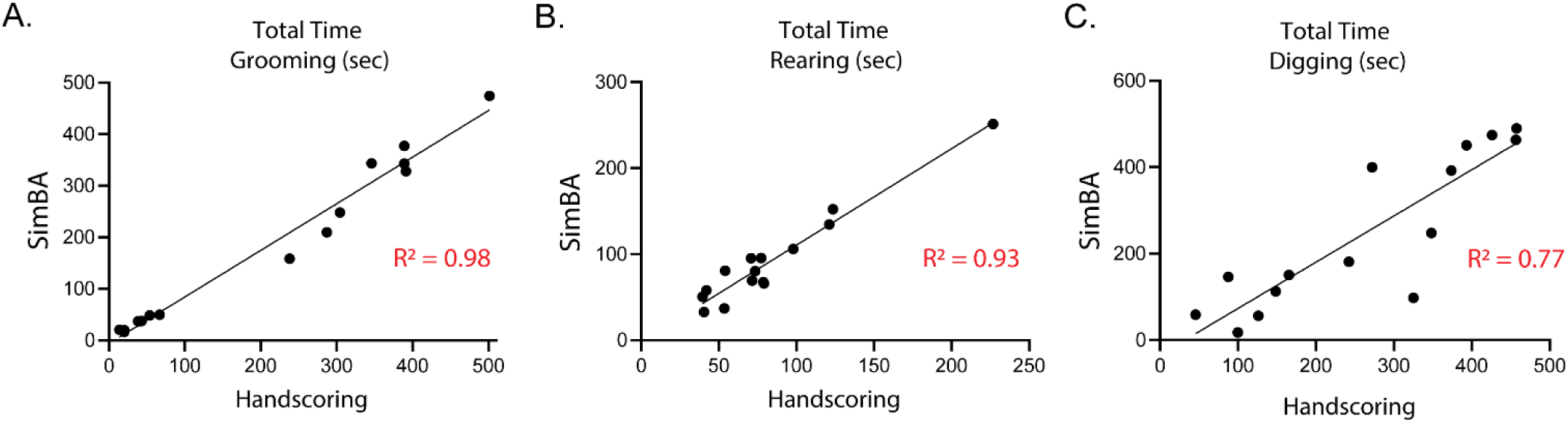
Traditional hand-scoring analysis and machine learning assisted analysis of behaviors are closely aligned. Linear Regression analysis of hand-score and SimBA analysis of time spent (A) Grooming (R^2^ = 0.98, p <0.0001]) (B) Rearing (R^2^ = 0.93, p <0.0001]) and (C) Digging (R^2^ = 0.77, p <0.0001]).

### Experiment 3

Next, we wanted to determine how long-term alcohol consumption impacts reactions to an ethologically relevant rodent predator task (Fig 4A). The looming disc task mimics an overhead predator and elicits active stress response behaviors, such as escape, that is associated with activation of paraventricular nucleus of the hypothalamus corticotropin releasing hormone (PVN^CRH^) neurons (Daviu et al., 2020).These neurons are altered in brain tissue of individuals with heavy alcohol use and are tightly involved in alcohol abuse and initiating the HPA-axis stress response (Sivukhina et al., 2006; Stephens and Wand, 2012). We compared the proportion of active escape, freezing, or neither escape nor freezing responses. In males, we found no significantly different behavioral responses between water and alcohol exposed mice (Fig 4B:Chi-Square test [X2 (1, N = 100) = 4.00, p = 0.135]). However, female alcohol exposed mice did differ in behavioral responses to this task compared to water controls (Fig 4C:Chi-Square test [X2 (1, N = 99) = 10.45, p = 0.005]). In particular, active escape reactions were seen more in alcohol mice (63% in alcohol vs 38% in water mice), while neither escape nor freezing reactions were more prevalent in water mice (12% vs 40% in water mice).

**Figure 4:**
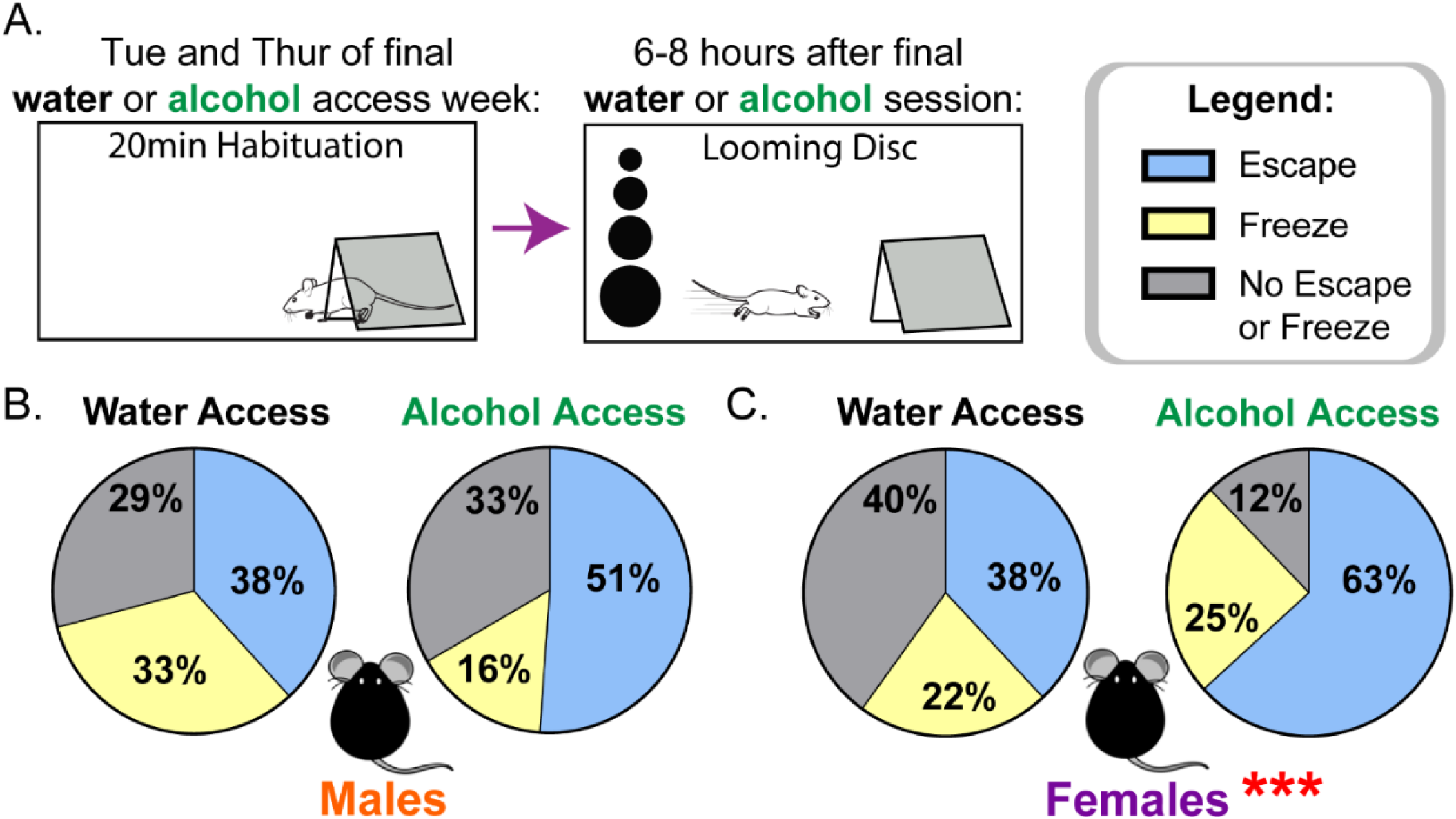
Female alcohol access mice in acute withdrawal escape more often to the looming disc than water access mice, but male alcohol and water mice do not differ in the behavioral reactions to the task. (A) Experimental design. (B) Proportion of Escape (Blue), Freeze (Yellow), and No escape or Freeze (Grey) in male water and alcohol exposed mice (Chi-Square test [X2 (1, N = 100) = 4.00, p = 0.135]). (C) Proportion of Escape (Blue), Freeze (Yellow), and No escape or Freeze (Grey) in female water and alcohol exposed mice (Chi-Square test [X2 (1, N = 99) = 10.45, p = 0.005]).

### Experiment 4

To determine if alcohol-exposed mice exhibited altered behavior in a more complex predator task that has been used to study the neural substrates of fear behavior (Amir et al., 2015; Choi and Kim, 2010), we examined foraging performance in the presence of a robogator threat during withdrawal. Beginning 7 days after the final drinking session, mice were given a 30-minute habituation session, 4 days of baseline training for pellet foraging, and a final Robogator test (Fig 5A). Mice showed no differences in multiple parameters of the habituation session, as depicted in the heat map for the first 5 minutes in the entire arena (Suppl fig 2A). Interestingly, alcohol access mice showed an overall higher latency to reach the pellet in the first training day (Fig 5B, Main effect of alcohol, [F(1, 16) = 7.017, p = 0.018]). Importantly, mice learned the foraging task and behaved equally by training day 4 before being introduced to the Robogator test the following day (Fig 5B), suggesting alcohol mice had higher anxiety-like behaviors in the beginning, but the differences in the test were likely not driven by differences in learning the task overall.

**Figure 5:**
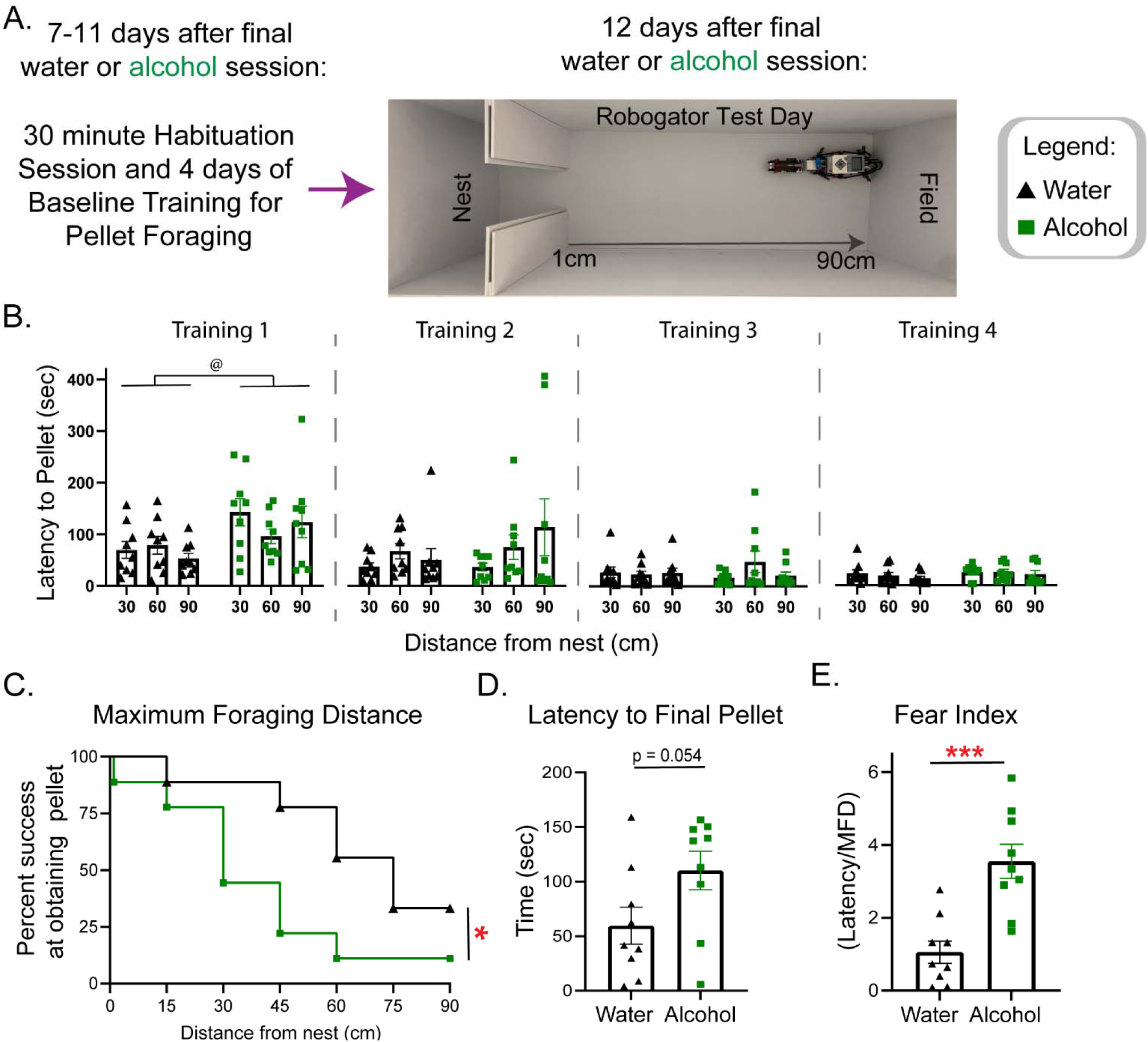
Alcohol access male mice in protracted withdrawal develop a more hesitant approach to foraging in the robogator task than water access mice. (A) Experimental design for the robogator task. (B) Latency to pellet for the 4 training sessions before test day, alcohol mice were slower overall in the first training day (RM 2-way ANOVA, Alcohol:p = 0.018). (C) Maximum Foraging Distance shown as percent success at obtaining the final pellet at each distance from nest, water access mice achieved farther distances in the presence of the gator than alcohol access mice (Log-rank (Mantel-Cox) test p = 0.0463). (D) Latency to final pellet (unpaired t-test p=0.0543). (E) Fear index shown as latency over maximum foraging distance; alcohol access mice had higher fear index (unpaired t-test p=0.0004).

A survival analysis was used to determine if there was a difference between groups in the Maximum Foraging Distance (MFD). We found a significant difference between water and alcohol mice, with water mice achieving further distances from the nest (Fig 5C, Log-rank (Mantel-Cox) test [X^2^(1, N = 18) = 3.970 p = 0.0463]). Although we found no differences in the latency to the final pellet (Fig 5D, [t(16)=2.077, p=0.0543]), when we normalized for the difference in the MFD from which each pellet was obtained by using the fear index (distance/latency), we found that alcohol mice scored significantly higher in this index than water mice (Fig 5E, [t(16)=4.462, p=0.0004]). Interestingly, we found no differences in the only shared trial by all mice, the 30cm first trial. The groups did not differ in the time it took to retrieve the pellet, the total distance travelled, number of nest entries, or percentage of time spent in the nest area (Supplementary fig 2B-E). Overall, a history of alcohol consumption in male mice increased the latency of foraging and anxiety-like behaviors in the presence of a predator-like threat.

## Discussion

In this study, we find that six weeks of two-bottle choice alcohol consumption impacts mouse behavioral responses to a home cage, open field, looming disc, and robogator predator test. While traditional anxiety/stress tests have been valuable in understanding alcohol induced alterations to negative affective behavior, stressors can induce different behavioral responses depending on factors such as stressor intensity/demands and task arousal. For instance, an open field test can induce anxiety-like behaviors due to moderately high arousal conditions from a bright light and novel environment (Roth and Katz, 1979). However, a stressor like the looming disc and robogator task, which pushes behavioral choices in a novel environment and in the presence of a predator-like threat, can induce a different arousal level and behavioral repertoire than the open field test. The responses to differing stressors may also be sex-specific, with some suggesting that female responses are biased towards stress-induced hyperarousal (Bangasser et al., 2018; Kudielka and Kirschbaum, 2005; Palanza and Parmigiani, 2017). Understanding how a history of voluntary alcohol consumption shifts behavioral responses to a variety of tasks during forced abstinence can lead to a wider understanding on alcohol induced changes to organisms and the brain.

To explore behavioral responses in a wide variety of tasks after long term voluntary alcohol consumption, we first assessed the two-bottle choice paradigm (Fig 1A). Male and female mice show high intake and preference for alcohol, but females display higher drinking patterns (Fig 1B-D). In a 4hr binge session, most mice in experiment 1 reached over 80mg/dL B.E.C.s, indicative of binge-like drinking behavior (Fig 1E). Interestingly, although few mice within an experimental cohort drink very low/large amounts of alcohol in an entire 24hr period, many of these parameters show subject variability in both males and females, leading to difficulties in examining behavioral correlations with alcohol intake. Thus, future experiments could seek to determine how differences in drinking patterns affect stress coping. As is often done with other models, it may be important for chronic voluntary drinking paradigms to take more frequent readings of B.E.C. and/or intake to characterize drinking more diligently within a 24hr period, similarly to what is done with non-human primates (Grant and Bennett, 2003), and what has been done to study individual differences in voluntary drinking in rodents (Heilig et al., 2019; Leeman et al., 2010; Spoelder et al., 2015). Overall, due to differences in baseline drinking patterns between males and females, this two-bottle choice access paradigm may not be conducive to investigating sex differences, and thus experiments 2 and 3 were analyzed separately.

Experiment 2 explored natural spontaneous behavior patterns in water and alcohol exposed mice following an acute stressor or in a naïve, no-stress, context. A recent investigation using a simple, low-anxiety inducing home-cage test showed specific spontaneous behavioral patterns in mice following acute stress. Interestingly, rearing and grooming in particular were driven by optogenetic manipulations of PVN^CRH^ neurons, which initiate the HPA-axis response and are involved in stress related behaviors (Füzesi et al., 2016). In rodents, rearing behaviors have been associated with increased exploration and vigilance and grooming is thought to be a coping behavior, though these behaviors can carry different meanings depending on the stressor type and intensity (Dielenberg et al., 2001; Fentress, 1988; Fernández-Teruel and Estanislau, 2016; Hermes et al., 2009; Sturman et al., 2018; Windle et al., 1997). In this low stress environment, we found that withdrawal from alcohol, in the absence of an acute stressor, shifts spontaneous behaviors similarly to a preceding stressor in males (Füzesi et al., 2016), in particular increasing the time spent rearing and reducing time spent engaging in digging behaviors (Figure 2D-E; I-J). While we did not find an increase in grooming behaviors, a correlation between total alcohol intake and time spent grooming was trending to significance (data not shown; R^2^ = 0.26, p = 0.08). Ongoing experiments using the same alcohol exposure paradigm and home cage test will continue to add to this data set.

While much more needs to be understood in relation to what alterations to spontaneous behaviors signify, it is interesting that acute withdrawal alone impacts natural home-cage behavior patterns in male mice. Particularly in regard to withdrawal from alcohol, this test in the absence of an immediately preceding stressor can prove highly useful, as traditional stress/anxiety tests encourage an anxiety-like state and can confound baseline, stress-induced withdrawal behaviors (Kliethermes, 2005). The home-cage test aims to reduce environmental factors that can increase variability and stress/anxiety by using the etho-experimental analysis of spontaneous animal behaviors in a low stress context. Additionally, these behaviors are sensitive to preceding stimuli, making the test valuable for measuring ‘stress coping’ in a natural, baseline environment (Füzesi et al., 2016; Lezak et al., 2017; Richter, 2020; Rojas-Carvajal et al., 2021). While the home cage test shows potential for the field, the analysis of these behaviors is extremely time consuming using traditional hand-scoring methods. To this end, we found that machine learning based analysis serves as a robust and reproducible alternative to hand-scoring that markedly reduces analysis time (Fig 3).

Our results suggest that restraint stress drives similar behaviors in male and female mice regardless of their alcohol exposure history (Fig 2). Compared to foot shock stress (Füzesi et al., 2016), restraint stress produced a much larger increase in grooming behaviors and a reduction, not increase, in time spent rearing. However, both stressors drastically reduce digging time. These home cage experiments are not directly comparable however, as all mice in our study were single housed for 8 or more weeks due to the requirements of the alcohol drinking paradigm. Social isolation itself is a stressor, particularly in females (Hermes et al., 2009; Mumtaz et al., 2018; Senst et al., 2016), which should be considered for the experiments performed in this study. Further research into how different stressors drive spontaneous home cage behaviors, and whether pharmacological treatment with drugs can restore these behaviors after stress can inform the field on the validity of this task.

Our interests also lie in understanding behavioral responses to immediate/acute stressors following a history of alcohol, particularly in species relevant predator tasks. The animated overhead looming disc, an ethologically relevant stressor that simulates a rapidly approaching overhead predator, has been shown to induce defensive behaviors in mice across many labs (De Franceschi et al., 2016; Shang et al., 2018; Yilmaz and Meister, 2013). Interestingly, escape responses are sensitive to changes in the activity of PVN^CRH^ neurons. A natural increase in activity precedes the initiation of escape and optogenetic inhibition of these cells decreases escape (Daviu et al., 2020). These neurons are altered by alcohol and withdrawal and are linked to various aspects of AUD (Sivukhina et al., 2006; Stephens and Wand, 2012). At an acute 6-8hr withdrawal timepoint, female, but not male, alcohol-exposed mice showed an increased propensity for active escape responses in the looming disc task (Fig 4). This finding in female mice might indicate increased activity of PVN^CRH^ neurons that predispose mice with a history of alcohol use to initiate an active coping strategy in response to stressors. Alcohol-exposed and naïve female mice did not show differences in the home-cage or open field test, which may be due to distinct coping strategies to male mice that appear as stressor intensity increases.

In 8-12 days forced abstinence, we found that male alcohol exposed mice showed increased latency to forage in a brightly lit environment in both the absence (Fig 5C) and presence (Fig 5D-F) of a predator-like robogator threat compared to water controls. The robogator task is a foraging task where mice are exposed to a moving predator threat that ‘attacks’ while they forage for food pellets. This task has been shown to involve brain regions closely related to anxiety/stress related behaviors, and the anxiolytic drug diazepam has been shown to attenuate risk aversion in a similar moving predator task (Amir et al., 2015; Choi and Kim, 2010; Walters et al., 2019). The findings with the robogator mirror those in the open field test at the 24hr withdrawal timepoint, where alcohol-exposed mice showed increased latency to enter the center of the open field (Fig 2G-H). Male alcohol mice also failed to show a more active stress coping strategy in the looming disc task (Fig 4). This avoidance/hesitancy under stress might suggest an underlying, universal phenotype seen after alcohol exposure in male mice.

Collectively, we find that a history of alcohol consumption leads to task-sensitive changes in behavior in mice. Mice display different sets of behavioral patterns under varied stress conditions, which may reflect the relative threat level of the assay. For example, the increased rearing at the cost of digging behaviors in male mice in the home-cage may reflect increased arousal and vigilance in a familiar environment. In contrast, the hesitancy in foraging in a brightly lit open environment and the findings with the open field tie more closely to increased anxiety states in these mice under higher arousal conditions. Female mice may not differ in behavioral patterns in low arousal conditions but begin to show alterations as the stressor intensity increases. It will be vital to continue studying how females respond to stress during withdrawal, as most research into negative affect in animal models has been conducted in males, while human research shows sex differences in alcohol and stress interactions during negative affective states (Becker and Koob, 2016; Brady and Sonne, 1999; Hilderbrand and Lasek, 2018; Peltier et al., 2019). Interestingly, behavioral effects in traditional anxiety/stress tests are often not seen in female rodents, which might hint at a need for different tests, such as those that incur heightened arousal or force behavioral choices under stress (Henricks et al., 2017; Jury et al., 2017; Reilly et al., 2009). Alternatively, given that several of these tasks were performed at different points post drinking, it is possible that the length of forced abstinence may impact behavioral performance. It is important to continue studying voluntary drinking paradigms, as there may be distinct neural alterations in a voluntary drinking model compared to forced access models. Almost no drug has advanced past clinical trials for treating alcohol withdrawal symptoms in recent years, and using new behavioral tests and voluntary drinking paradigms may help expand our understanding of alcohol use and abuse (Heilig et al., 2019). Finding new and efficient treatments could encourage people with AUD to seek treatment earlier in the history of alcohol use, reduce withdrawal associated negative affective states and the severity of AUD (Finn and Crabbe, 1997; Glass et al., 2017).

Earlier intervention strategies can aid in harm reduction for overall better health outcomes (Charlet and Heinz, 2017).

---

**Supplementary Figure 1:**
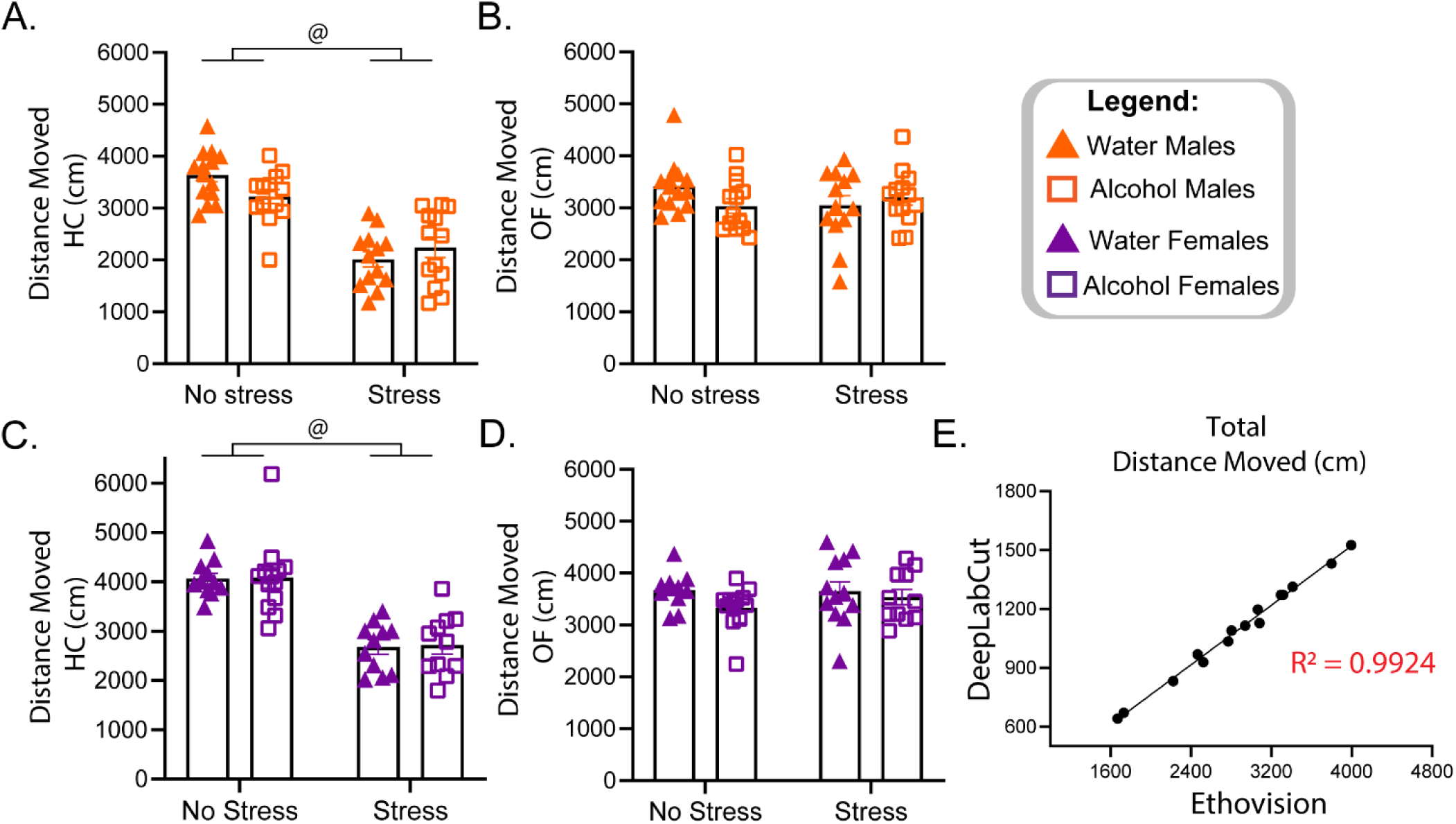
Locomotion effects in the home cage and open field test. (A) Stress male mice show a significant reduction in distance moved in the home cage test (Two-way ANOVA, Main effect of Stress Condition [F(1, 49) = 71.14, p<.001]) (B) No significant differences between male groups in the distance moved in the open field test. (C)Stress female mice show a significant reduction in distance moved in the home cage test (Two-way ANOVA, Main effect of Stress Condition [F(1, 42) = 63.33, p<.001]) (D)No significant differences between female groups in the distance moved in the open field test. (E) Comparison of Ethovision derived locomotion data to DeepLabCut derived data for the home cage test (simple linear regression:[F(1, 13) = 699.3, p <.0001]).

**Supplementary Figure 2:**
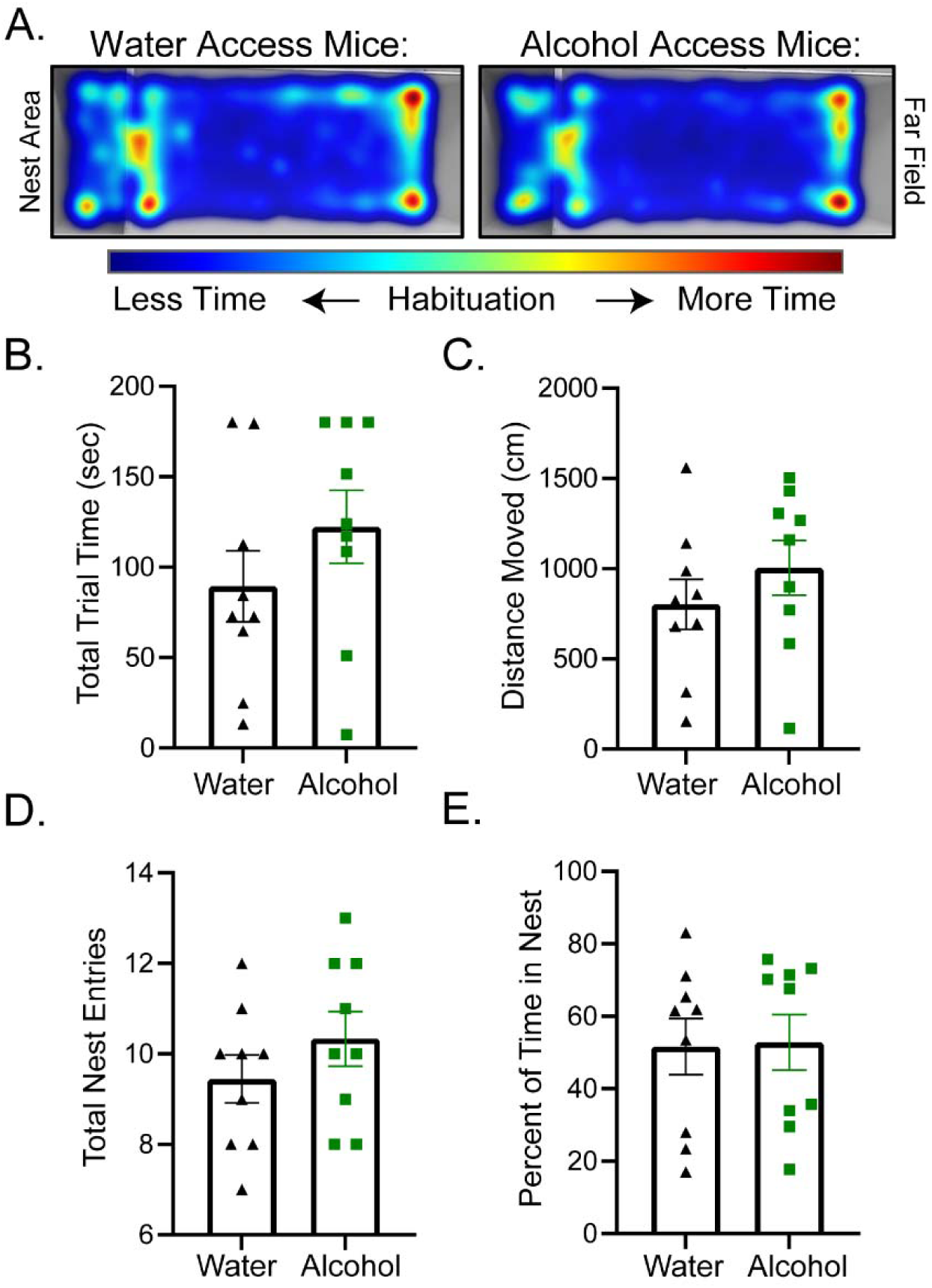
No differences in the robogator task between water and alcohol access mice in habituation or the first, shared, 30cm foraging trial. (A) Heat map depicting the first 5 minutes of the habituation session shows no differences between groups. Parameters for shared 30cm trial, with maximum trial time being 180 seconds:(B) Total trial time (C) Distance moved (D) Total nest entries (E) Percent of time spent in nest.

## Acknowledgements

This work was supported by grants from the Howard Hughes Medical Institute James H. Gilliam, Jr. Fellowship for Advanced Study (GT13514) and the National Institute of Health (U01AA020911, U24AA025475, P60AA011605). Special acknowledgements to Dr. Dipanwita Pati and Dr. Melanie Pina for their invaluable mentorship and feedback. Thanks to SciDraw, a free website where people donate their fantastic drawings for others to use, and BioRender.com, which together allowed me to create my figures.

## Contributions

S.N., J.S.B., and T.L.K. conceptualized experiments. S.N. performed behavioral experiments, data analysis, and manuscript writing. L.A.H. performed home cage and open field experiments in females. C.M.S assisted with behavioral experiments and behavioral analysis. M.C.B. aided with behavioral analysis. S.L.D. and H.L.H. assisted with drinking experiments. M.E.F. established DLC/SimBA in the lab and aided with analysis. K.M.B. assisted with behavioral experiments. All authors contributed to the editing of the manuscript.7

## Ethics Declarations

The authors declare no competing interests.

## Additional Information

N/A

## Notes

### Competing Interest Statement

The authors have declared no competing interest.

